# Filamentation activates bacterial NLR-like antiviral protein

**DOI:** 10.1101/2024.09.15.613165

**Authors:** Yiqun Wang, Yuqing Tian, Xu Yang, Feng Yu, Jianting Zheng

## Abstract

Bacterial antiviral STANDs (Avs) are evolutionarily related to the nucleotide-binding leucine-rich repeat containing receptors (NLRs) widely distributed in immune systems across animals and plants. *Ef*Avs5, an Avs type 5 protein from *Escherichia fergusonii*, contains an N-terminal SIR2 effector domain, a nucleotide-binding oligomerization domain (NOD) and a C-terminal sensor domain, conferring protection against diverse phage invasions. Despite the established roles of SIR2 and STAND in prokaryotic and eukaryotic immunity, the mechanism underlying their collaboration remains unclear. Here we present cryo-EM structures of *Ef*Avs5 filaments, elucidating the mechanisms of dimerization, filamentation, filament clustering, ATP binding and NAD^+^ hydrolysis, all of which are crucial for anti-phage defense. The SIR2 domains and NODs engage in the intra- and inter-dimer interaction to form an individual filament, while the outward C-terminal domains contribute to bundle formation. Filamentation potentially stabilizes the dimeric SIR2 configuration, thereby activating the NADase activity of *Ef*Avs5. *Ef*Avs5 is deficient in the ATPase activity, but elevated ATP concentrations can impede its NADase activity. Together, we uncover the filament assembly of Avs5 as a unique mechanism to switch enzyme activities and perform anti-phage defenses, emphasizing the conserved role of filamentation in immune signaling across diverse life forms.

**Keypoints:**

- *Ef*Avs5 depletes NAD^+^ for anti-phage defense.
- *Ef*Avs5 assembles as bundled filaments that can hydrolyze NAD^+^.
- The SIR2 domain and NOD collaborate to form an individual filament.
- The building block of the filament is *Ef*Avs5 dimer.
- The activity of the *Ef*Avs5 complex is regulated by ATP.

## Introduction

Bacteria are in constant conflict with bacteriophages and develop diverse anti-phage defense systems of varying complexity^1-3^. The signal transduction ATPases with numerous domains (STAND) superfamily of P-loop NTPases, particularly the nucleotide-binding leucine-rich repeat containing receptors (NLRs) exemplified by animal inflammasomes and plant resistosomes^4-6^, recognizes pathogens and triggers downstream inflammatory responses or apoptotic processes through oligomerization, representing a conserved immune strategy in eukaryotes. Recently, this mechanism has also been discovered in the innate immunity of bacteria, known as antiviral STANDs (Avs)^3^. Avs proteins have a characteristic tripartite domain architecture: a central NTPase (alternatively called nucleotide-binding oligomerization domain, NOD); an extended C-terminal sensor with superstructure-forming repeats; and a variable N-terminal effector typically involved in cell death. Previous studies have revealed that Avs1-3 and Avs4 are activated by the phage terminases and portals, respectively, leading to the formation of tetramers that activate their N-terminal nucleases for antiviral defense^4^.

Avs type 5 (Avs5) contains an N-terminal sirtuin (SIR2) effector and an unusual short C-terminal sensor domain, sometimes termed SIR2-STAND^3^. SIR2-mediated nicotinamide adenine dinucleotide hydrolase (NADase) activity represents a critical element in anti-phage immunity, triggering abortive infections via NAD^+^ degradation and halting phage propagation, as exemplified by pAgo, Thoeris, DSR2, and the SIR2-HerA system^7-10^. In the Thoeris system, the signaling molecule 1’’-3’ gcADPR activates ThsA, which subsequently forms filaments and depletes NAD^+^, ultimately triggering cell death and thereby preventing the spread of phage infection^11^. Another defense system DSR2 tetramerizes to form a supramolecular complex that specifically recognizes phage tail tube proteins, and leads to cellular NAD^+^ depletion^12^.

Despite reports on SIR2 and STAND roles in prokaryotic and eukaryotic immunity, the collaborative mechanism between them and the mechanisms underlying Avs5 activation and assembly remain obscure. Here, we present a biochemical and structural analysis of *Escherichia fergusonii* Avs5 (*Ef*Avs5), unveiling a unique mechanism of assembly-mediated activation by *Ef*Avs5. *Ef*Avs5 proteins form active filaments via SIR2 and NOD self-association, with sensor-mediated clustering enhancing their organization, allowing the NAD^+^ depletion during phage infection. Moreover, the inactive NOD of *Ef*Avs5 requires ATP binding for its defensive role, whereas high ATP concentration impedes its NADase activity.

## Result

### 1. *Ef*Avs5 consumes NAD^+^ to confer defense

Avs5 system is widely distributed across Gram-negative phyla, especially Pseudomonadota (Fig. S1). To define the molecular anti-phage defense mechanism of Avs5, we investigated a single protein Avs5 defense system from *E. fergusonii* phage-inducible chromosomal islands (PICI), named *Ef*Avs5 (Fig. 1A). *Ef*Avs5 is reported to defense against diverse *Escherichia coli, Salmonella* and *Klebsiella pneumoniae* phages^13^. We cultured *E. coli* cells expressing *Ef*Avs5 and found protection against T7 phages. The C terminal of *Ef*Avs5 is supposed to sense the phage invasion, while the SIR2 domain is suggested to be the effector depleting NAD^+^. Indeed, a single amino acid substitution at NADase active site (N141A) was sufficient to abolish phage defense by the *Ef*Avs5 system, confirming that the NADase activity is essential for defense. Similarly, a point mutation disrupting the NOD active site, K379A in walker A or D426A in walker B, as well as a deletion of the C-terminal sensor domain, also abolished defense (Fig. 1A).

**Fig. 1.**
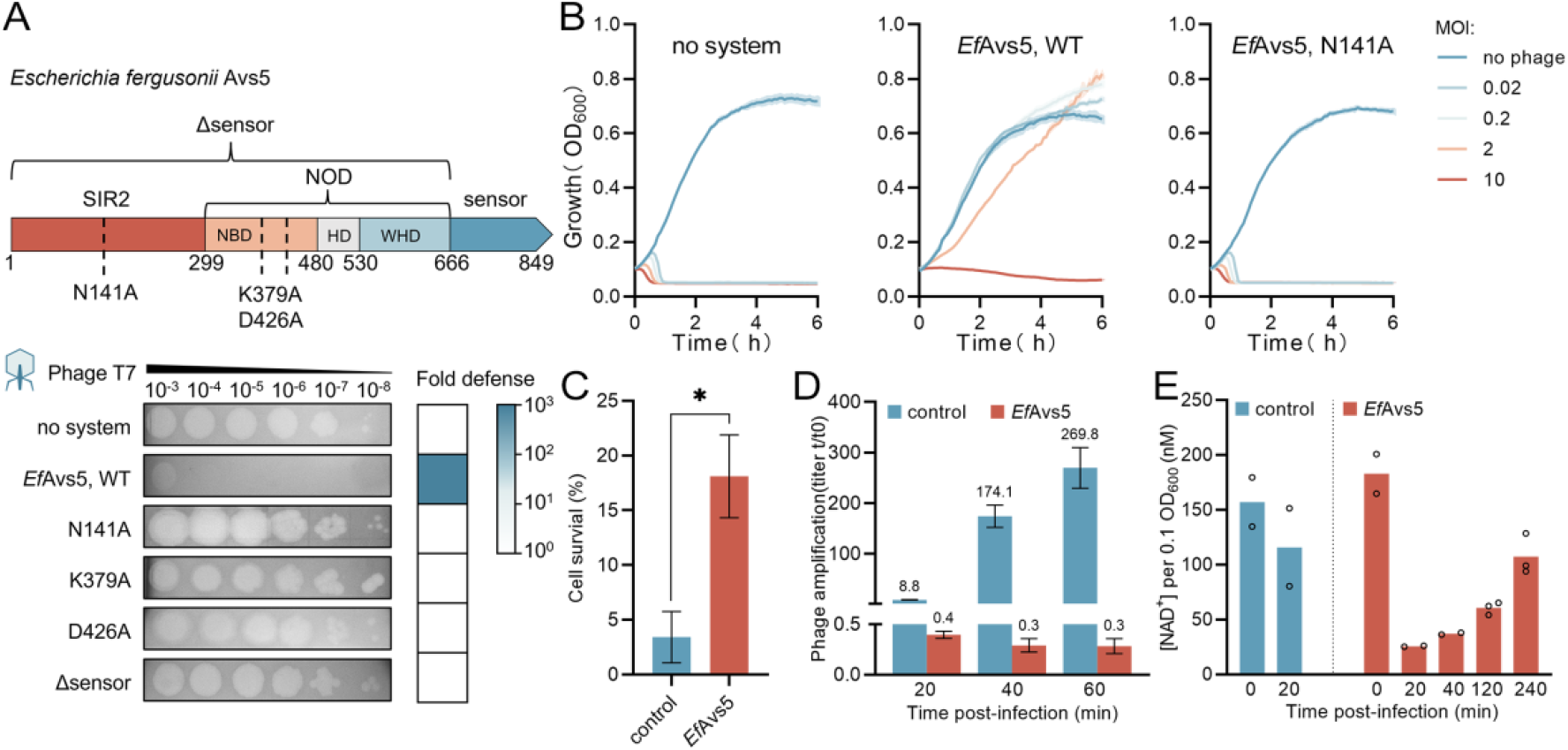
*Ef*Avs5 consumes NAD^+^ to confer defense. (**A**) Domain organization of the *Ef*Avs5 protein from *Escherichia fergusonii* and efficiency of plating of T7 phages infecting *E. coli* expressing *Ef*Avs5 or an empty vector. Data are representative images of n=3 biological replicates. Mutations used are indicated below the domain organization. NBD, nucleotide-binding domain; HD, helical domain; WHD, wing-helical domain; NOD, nucleotide-binding oligomerization domain. (**B**) Infection time courses for liquid cultures of *E. coli*, with *Ef*Avs5, *Ef*Avs5 N141A or empty vector, infected at different multiplicities of infection (MOI) of phage T7. Data represent the mean of three replicates and shaded regions represent the standard error of the mean (SEM). (**C**) Survival of *E. coli* cells expressing either the *Ef*Avs5 or empty vector as a control infected at an MOI of 5 with T7 for 20 minutes. Data represent the mean of three replicates and error bars represent the SEM. (**D**) Titer of T7 phage propagated on *E. coli* cells expressing either the *Ef*Avs5 or empty vector as a control infected at an MOI of 0.1 with T7. Data represent the titer of T7 measured in PFU/mL after indicated time from initial infection, divided by the original phage titer prior to infection. Data represent the mean of three replicates and error bars represent the SEM. (**E**) Measurement of NAD^+^ concentration in *E. coli* expressing *Ef*Avs5 or empty vector as a control at the indicated time points after infection with phage T7 at an MOI of 2, normalized to an OD_600_ of 0.1. The control group underwent near-complete lysis after 20 minutes of infection, precluding the quantification of NAD^+^ content. Data represent the mean of two or three replicates and error bars represent the SEM.

Growth curves of *E. coli* cultures in the presence of *Ef*Avs5 kept growing when subjected to the low (0.2) or medium (2) multiplicity of infection (MOI) in contrast to the cells expressing N141A mutant or lacking *Ef*Avs5 (Fig. 1B). We measured the survival rate of cells infected with T7 and found that *Ef*Avs5 increased cell survival after the first infection cycle (about 20 minutes) (Fig. 1C). Further analysis showed that the phages amplified rapidly in the infected cells lacking *Ef*Avs5, but not in the presence of *Ef*Avs5 (Fig. 1D). To investigate the effector’s activity during immunity, we detected the cellular NAD^+^ content after infection with T7 phage. When subjected to the T7 phage infection, compared to cells lacking *Ef*Avs5, the concentration of NAD^+^ in the *Ef*Avs5-expressing cells was declining rapidly after the first infection cycle, and began to rise in the second infection cycle (Fig. 1E). These results suggest that *Ef*Avs5 consumes NAD^+^ to effectively prevent phage amplification and leads to the population-level protection.

### 2. Architecture of *Ef*Avs5 filament assembly

To biochemically characterize the Avs5 system, we purified *Ef*Avs5 protein overexpressed in *E. coli* BL21(DE3). The freshly isolated *Ef*Avs5 exists as soluble monomer (Fig. S2). Cryo-EM imaging of the monomer *Ef*Avs5 revealed small particles. We tried to resolve the structure but only obtained two-dimensional (2D) classes in poor resolution, possibly due to the flexibility (Fig. S3). However, the purified *Ef*Avs5 becomes visibly cloudy after stored at 4°C or -80°C for two weeks. Negative staining analysis of the suspension of *Ef*Avs5 confirmed that a significant fraction of the particles (>50%) exhibited bundled 8-nm-wide fibrous assembly, and some of them extending up to ∼400 nm in length (Fig. 2A). Bundled fibrils can be connected parallelly or end-to-end and extend up to several micrometers. Since previous studies show that the activated SIR2 or Toll/interleukin-1 receptor (TIR) can assemble into helical filaments in Thoeris defense system, TIR-STING and TIR-SAVED effector, respectively^11,14,15^, we quantified the NADase activity of *Ef*Avs5. The freshly purified *Ef*Avs5 was deficient in the NADase activity, but the filamentous *Ef*Avs5 displayed robust NADase activity (Fig. 2B).

**Fig2.**
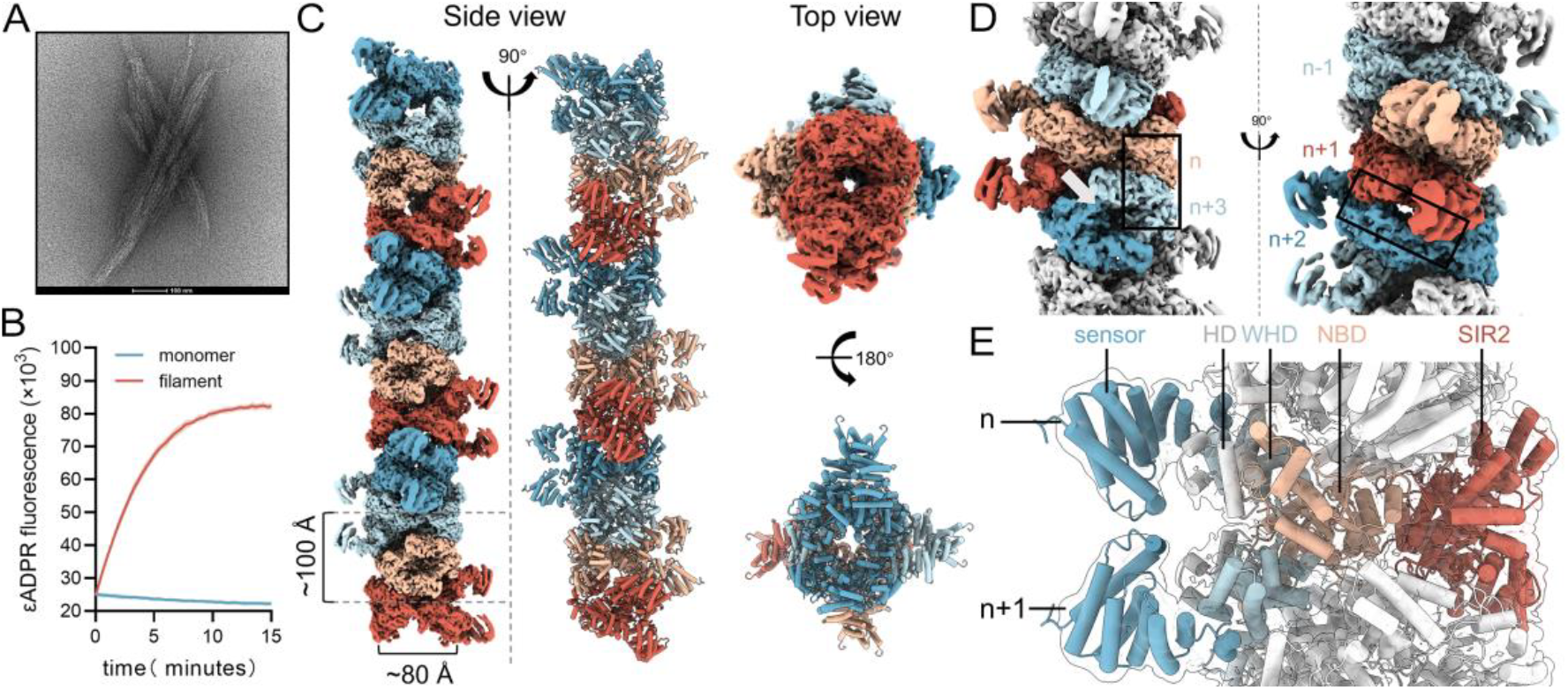
Cryo-EM structure of the *Ef*Avs5 filament. **(A)** Representative negative-stain micrograph images of *Ef*Avs5. The scale bar represents 100 nm. **(B)** *Ef*Avs5’s NADase activity is activated by filamentation. After the addition of ε-NAD^+^, total fluorescence was measured over time. Data represent the mean of three replicates and shaded regions represent the SEM. (**C**) Top and side views of the density map and the atomic model of *Ef*Avs5 filament. (**D**) Detailed view of *Ef*Avs5 filament to show the subunit interfaces. Two interfaces are indicated with boxes, and the NAD^+^ active site is indicated with a grey arrow. (**E**) The core of *Ef*Avs5 filament is composed of the SIR2 domain and the NOD.

To gain more insight into the filament formation, we determined the cryo-electron microscopy (cryo-EM) structure of the active *Ef*Avs5. The final cryo-EM map was reconstructed using a total of 228,053 single particles and refined to a nominal resolution of 3.4 Å, with approximately 2.8 Å at the center of filament and approximately 6.8 Å at the peripheral sensor lobes (Fig. 2C, Fig. S4 and Table S1). The model was built with eight *Ef*Avs5 molecules assembled into a helical filament, and extra densities suggest unlimited entries of both ends. The repeating building block of the filament is *Ef*Avs5 dimer with a C_2_ symmetry arranged into a regular helical structure with a helical twist of 84.531° and a helical rise of 48.600 Å (Fig. 2C). The filament is held together by 2 sets of inter-subunit interactions, one between subunit n and n+3 and the other between subunit n+1 and n+2 (Fig. 2D). The *Ef*Avs5 proteins fold into five domains: a catalytic SIR2 domain, a core NOD, which comprises the nucleotide-binding domain (NBD), helical domain (HD) and wing-helical domain (WHD) and the C-terminal sensor domain (Fig. 1A and Fig. 2E). The core of the helix is composed of the SIR2 and the NOD domain, whereas the C-terminal domain is pointing away and not involved in the helical contacts (Fig. 2E).

### 3. The dimerization of *Ef*Avs5

In the *Ef*Avs5 filament, the individual dimeric unit adopts a C_2_ symmetric conformation with a tight interface (2329.2 Å^2^ of buried surface area) (Fig. 3A). Within each *Ef*Avs5 dimer unit, abundant contacts are observed between the WHD of one subunit and the NBD of its adjacent subunit, encompassing salt bridges and hydrogen bonds, revealing an oligomerization pattern distinct from other characterized active STAND ATPases, which primarily oligomerize through NBD-NBD and WHD-WHD interactions (Fig. S5). The NBD of the n subunit interacts with the WHD of the n+1 subunit, comprising the salt bridge of the K461(n)-E589(n+1), K473(n)-E580(n+1), and the hydrogen bonds of the T474(n)-Y583/R570(n+1) (Fig. 3B). Meanwhile, the WHD of the n subunit interacts with the NBD of the n+1 subunit using slightly different residues, comprising the specific contacts of the E580(n)-H362(n+1), and the hydrogen bonds of the Y583(n)-T474(n+1). The dimeric interface is expanded by additional hydrogen bonds between its N-terminal SIR2 domains (Fig. 3C). To probe the role of the dimer formation in *Ef*Avs5 activity, we generated mutants of the dimeric interface. The substitutions of Y241 and T262 in SIR2 domain, H362A and K461 in NOD abolished *Ef*Avs5 system antiviral defense, indicating that dimer formation is obligatory for *Ef*Avs5 anti-phage activity *in vivo* (Fig. 3D).

**Fig3.**
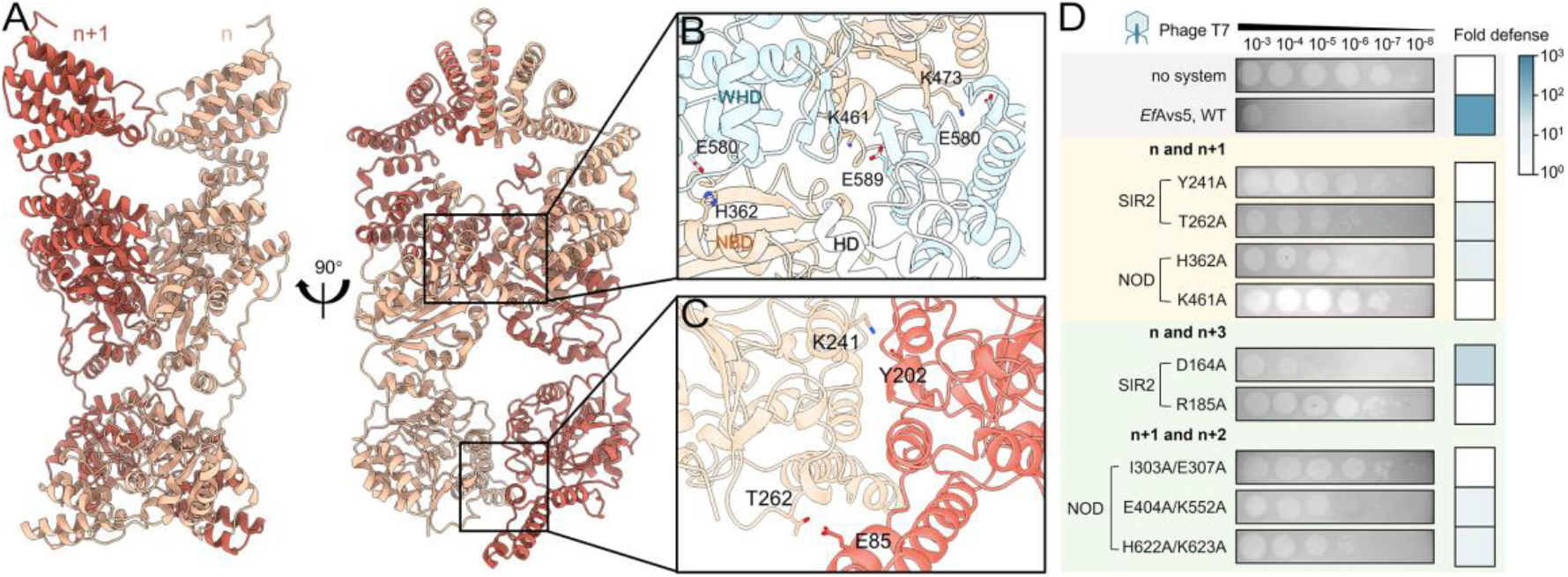
The mechanism of *Ef*Avs5 dimerization. (**A**) The structure of the *Ef*Avs5 dimer. (**B**) Details of the NOD-based dimeric interfaces. (**C**) Details of the SIR2-based dimeric interfaces. (**D**) Phage plaque assay of *E. coli* expressing *Ef*Avs5 variants with mutations at the interfaces. The yellow background highlights the intra-dimer interactions, while the green background indicates the inter-dimer interactions. Data are representative images of n=3 biological replicates.

### 4. SIR2- and NOD-dependent filamentation

The *Ef*Avs5 structure features both SIR2 and NOD engaging in intra- and inter-dimer interactions, resulting in two spirals: one dominated by SIR2, the other by NOD (Fig. 4A, B and E). The SIR2 domain within *Ef*Avs5 system exhibits a typical two-domain fold (Fig. 4D), resembling DSR2(PDB ID: 8WYB, rmsd of 1.160 Å for 74 aligned Cα atoms) and ThsA (PDB ID: 8BTO, rmsd of 1.247 Å for 84 aligned Cα atoms)^11,12^, except for a longer α3 creating a broader substrate channel (Fig. S6). The large domain (LD) has a Rossmann-fold core that is conserved among all the SIR2 proteins, while the small domain (SD) lacks the three-stranded zinc-binding motif compared to human Sirt5^16^.

**Fig 4.**
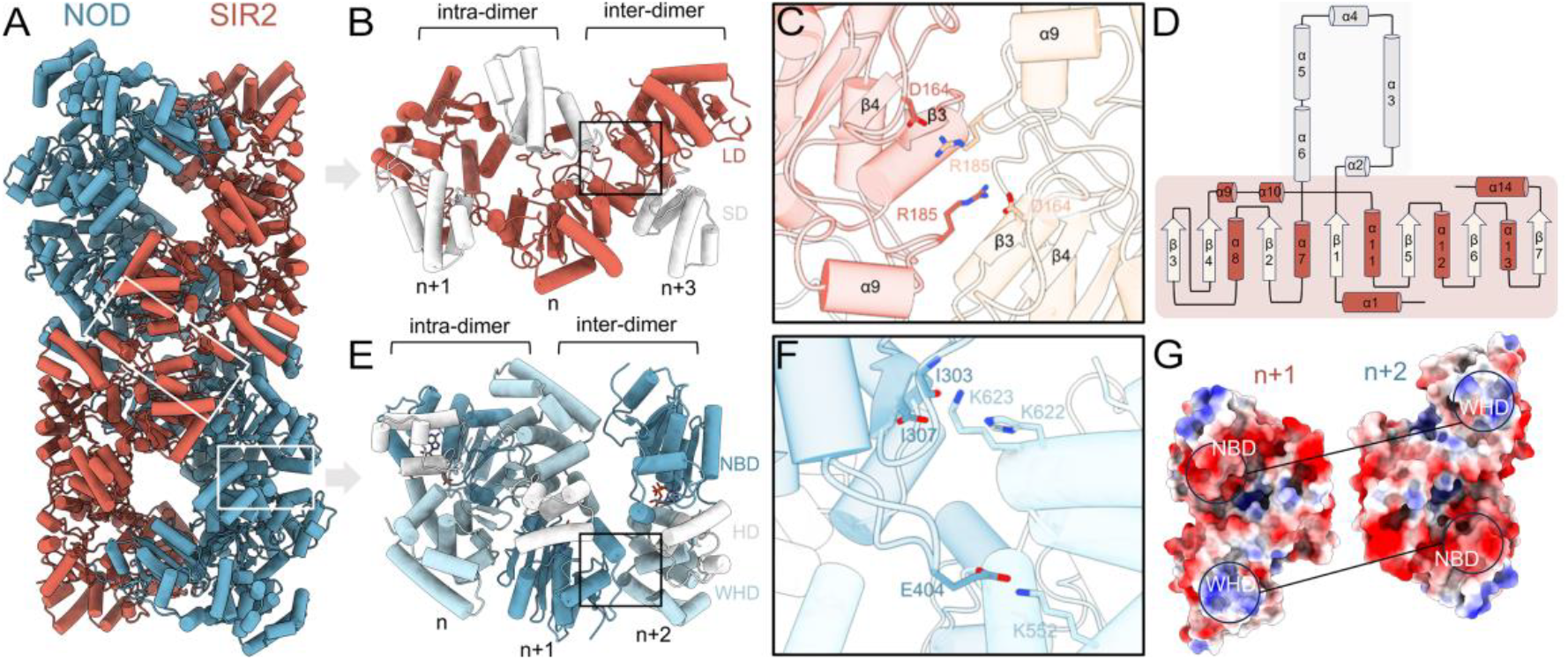
SIR2- and NOD-dependent filamentation. (**A**)Two spirals dominated by NOD (blue) and SIR2 domains (red), respectively. The C-terminal domains of *Ef*Avs5 are hided. The inter-dimer interactions are indicated by white boxes. (**B**) The intra-dimer and inter-dimer interface of SIR2 domain of the n+1, n, n+3 subunits. LD, large domain; SD, small domain. (**C**) Detailed inter-dimer interface of SIR2 domain. The n, n+3 subunits of *Ef*Avs5 was colored red and orange, respectively. (**D**) SIR2 topology diagrams of *Ef*Avs5. The small domain is shaded grey, and the large domain is shaded red. (**E**) The intra-dimer and inter-dimer interface of NOD domain of the n, n+1, n+2 subunits. (**F**) Detailed inter-dimer interface of NOD domain. The NBD of n+1 subunit, and the WHD of n+2 subunit of *Ef*Avs5 were colored dark and light blue, respectively. (**G**) The electrostatic potential inter-dimer interface of two adjacent NOD domains.

The interface between SIR2 dimers is non-parallel to the helical axis (Fig. 4A), burying ∼1218.8 Å^2^ of surface area (Fig. 4B). SIR2 α4, α10 and α12 helices contribute to the intra-dimer interactions (Fig. 3C), whereas the loop connecting β3 and β4 forms polar interactions and hydrogen bonds with the loop between β4 and α9 of the adjacent SIR2, thereby constituting the inter-dimer interactions and facilitating the assembly of *Ef*Avs5 filaments (Fig. 4C). Interestingly, despite the similarity of the structure, the oligomerization mechanism of *Ef*Avs5-SIR2 is different from SIR2-HerA, ThsA and DSR2 (Fig. S7). Identical to the intra-dimer interaction, the inter-dimer interactions of NOD also mainly include the NBD-WHD contacts, but adopt a different mechanism (Fig. 4E). The NBD interface of the n+1 molecule is predominantly negatively charged (I303, E307, E404), complementing the positively charged interface of WHD domain of n+2 molecule (K552, H622, K623) (Fig. 4F and G).

To probe the role of the filament formation in *Ef*Avs5 activity, we generated mutants D164A and R185A of the interface between n and n+3 subunits, and double mutants I303A/E307A, E404A/K552A and H622A/K623A of the interface between n+1 and n+2 subunits. These substitutions abolished *Ef*Avs5 system antiviral defense, indicating that both SIR2-based and NOD-based filament formation is obligatory for *Ef*Avs5 anti-phage activity *in vivo* (Fig. 3D).

### 5. NADase catalytic pocket

As expected, modeling of NAD^+^ within the active site in the cryo-EM map of *Ef*Avs5 was not feasible due to degradation by the active SIR2. Superimposing the SIR2 domain of *Ef*Avs5 onto both the active and inactive conformations of ThsA reveals a comparable active pocket, situated within the SIR2 dimer (Fig. 5A and Fig. 2D). Dimerization likely contributes to the stabilization of the active pocket. In the case of ThsA and DSR2, the position of the loop above the active pocket decides to block the NAD^+^ access to the active site or enable NAD^+^ binding, thus control the NADase activity of SIR2^11,17^. However, in the *Ef*Avs5, the longer α3 pull this loop far away the active pocket (Fig. 5A and Fig. S6), indicating a different activation mechanism adopted by *Ef*Avs5. In the *Ef*Avs5-NAD^+^ predicted model, the A41, T140, N141, Y142, N180, Y202, S233 in the pocket interact with NAD^+^, creating an ideal condition for the reaction (Fig. 5B). The Y202 is also involved in SIR2-based dimerization. These residues are highly conserved in the bacterial SIR2 proteins, except the substitution of an Asn residue with His at residue 180 in *Ef*Avs5 (Fig. S8).

**Fig 5.**
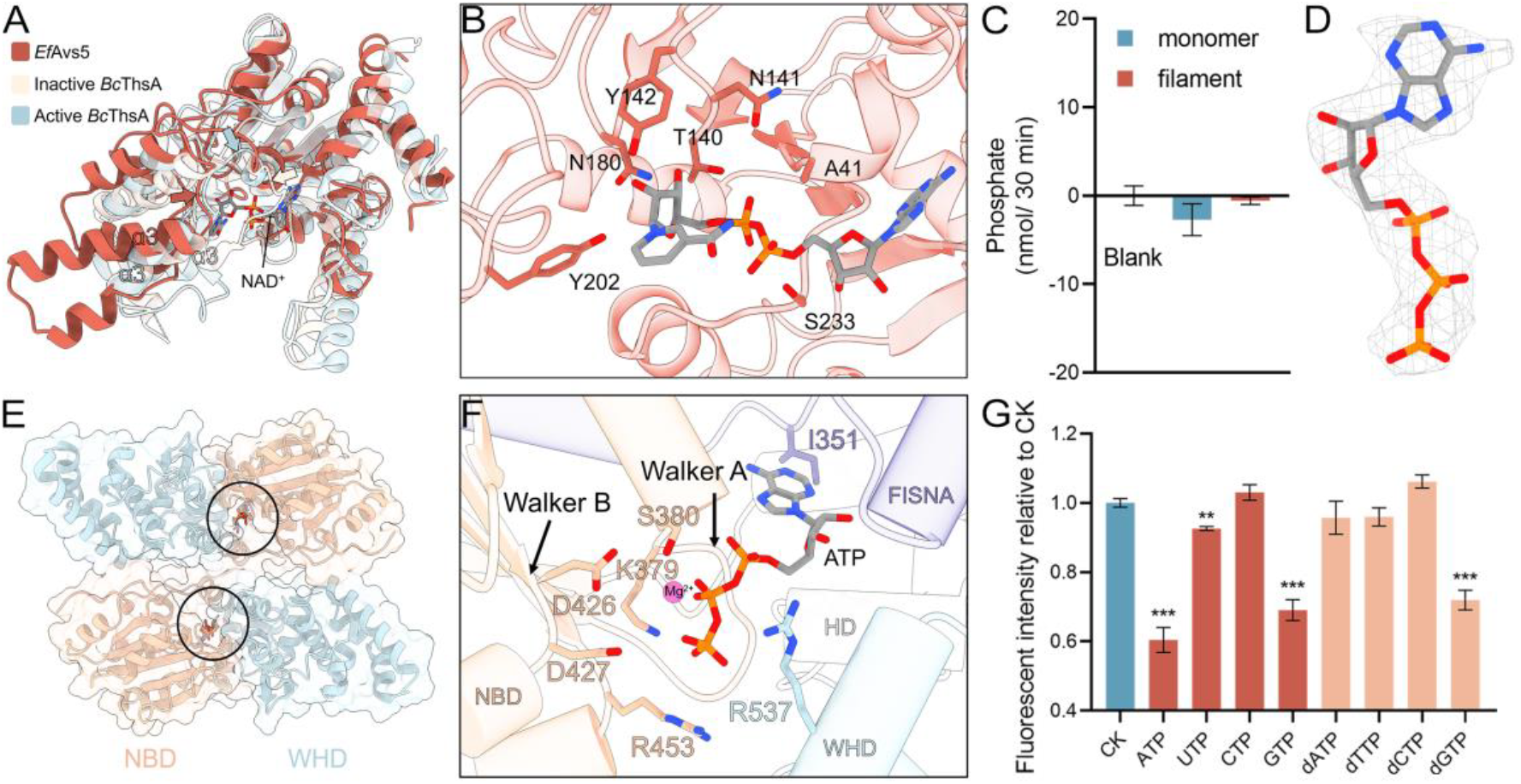
The NAD^+^ hydrolyzing pocket and ATP binding pocket of *Ef*Avs5. (**A**) Overlay of SIR2 domain (red) of *Ef*Avs5 with the inactive (6LHX, light orange) and activated (light blue) *Bc*ThsA SIR2 domains. NAD^+^ from the activated ThsA (1ICI) is shown. The loops above the NAD^+^ pocket of *Ef*Avs5 and ThsA SIR2 are indicated by arrows colored as in the color scheme. (**B**) Residues of *Ef*Avs5 interacting with NAD^+^ molecule highlighted in sticks in the predicted *Ef*Avs5-NAD^+^ binding model. (**C**)ATPase activity assay of the monomer or the filament *Ef*Avs5. Data represent the mean of three replicates and error bars represent the SEM. (**D**) Cryo-EM map of ATP. (**E**) The binding pocket of ATP in the NOD dimer. (**F**) Residues of *Ef*Avs5 coordinating ATP molecule highlighted in sticks. The highly conserved Walker A motif and Walker B motif were indicated by arrows. (**G**) Effects of NTPs and dNTPs (5 mM) on NADase activity of the *Ef*Avs5. CK: no NTPs/dNTPs added. Data represent the mean of three replicates and error bars represent the SEM.

### 6. ATP inhibits the NADase activity of the *Ef*Avs5 system

The NOD of *Ef*Avs5 is an AAA^+^ ATPase, belongs to a novel STAND NTPase 3 family found in bacterial conflict systems and in metazoan TRADD-N associated counter-invader proteins^18^. The structural comparison in Dali server^19^ suggested that *Ef*Avs5’s NOD is most similar to the plant NLR RPP1(PDB ID: 7CRC, rmsd of 1.160 Å for 56 aligned Cα atoms) and cell death protein 4 in *Caenorhabditis elegans* (PDB ID: 4M9S, rmsd of 1.239 Å for 45 aligned Cα atoms). Notably, we found that *Ef*Avs5 is an inactive ATPase either in the monomer or filament assembly (Fig. 5C). Consistently, the cryo-EM map allows modelling of ATP molecules in each NOD active site (Fig. 5D), although we did not add additional ATP during the sample preparation. In the structure, ATP molecules are buried in the active pocket between NBD and WHD (Fig. 5E), with an adjacent magnesium ion coordinated by the canonical Walker A and B motif (Fig. 5F). The K379 and S380 in Walker A form hydrogen bonds with the oxygens of the phosphate groups, determining the orientation of ATP. Meanwhile, the acidic D426 in Walker B binds to the Mg^2+^. The recognition of the γ-phosphate group of ATP in the ZAR1 resistosome and Apaf-1 apoptosome is facilitated by an arginine residue in the conserved ‘TT/SR’ motif, which is crucial for their activation and preserved as ‘TTR’ and R453 in *Ef*Avs5^20,21^. The I351 forms hydrogen bonds with the N6 atoms of the adenine ring (Fig. 5F). The region spanning residues 300-366, sometimes described as the fish-specific NACHT-associated (FISNA) domain, resembles the active state of NLRP3 (PDB ID:8EJ4), triggering conformational alterations and oligomerization^22^. The accurate positioning of ATP is crucial for the functionality of the *Ef*Avs5 system, as evidenced by the phage plaque assays with WalkerA/B mutants (Fig. 1A). The deficiency of ATP hydrolysis probably results from its WHD domain’s unique conformations near the active site, particularly the hydrogen bonding between R537 and the ATP phosphate, contrasting with other active STAND ATPases (Fig. 5F and Fig. S9).

Based on the role of ATPase in sustaining *Ef*Avs5’s activity *in vivo*, we investigated the potential impact of ATP on NAD^+^ hydrolysis. Our findings revealed that low ATP concentrations (0.05 and 0.5 mM) had negligible effects, whereas 5 mM ATP significantly reduced *Ef*Avs5’s NADase activity by 40% (Fig. S10). Notably, GTP and dGTP also effectively inhibited NADase activity, while other NTPs/dNTPs showed minimal effects, except for a slight inhibition by UTP (Fig. 5G). These results indicate that specific NTP/dNTP and its concentration can inhibit *Ef*Avs5’s NADase activities.

### 7. Filamentous cluster formation mediates *Ef*Avs5 anti-phage activity

*Ef*Avs5 filaments are organized into higher-order structures, forming bundles or three-dimensional networks (Fig. 2A). Although density maps of the filament bundle were not available, interactions between filaments were observed during 2D classification in cryo-EM analysis (Fig. 6A), in which the *Ef*Avs5 exhibits an end-to-end linkage between dimers from adjacent filaments (Fig. 6B). Notably, the C-terminal sensor possesses a predominantly positive charge, which complements the negatively charged interface of the N-terminal SIR2 domain residing in adjacent filaments (Figure 6C). These interactions would enable the *Ef*Avs5 filament to extend in four distinct directions, thereby promoting the formation of parallel bundles (Figure 6D). The C-terminus of *Ef*Avs5 does not participate in the formation of the individual filament but may play a crucial role in filament clustering. Deletion of the entire C-terminal sensor abolished *Ef*Avs5 activity (Fig. 1A). Further analysis confirmed that deletion of C-terminal six positively charged amino acids similarly abolished *Ef*Avs5’s resistance to phage invasion, as well as the mutations of D79A, E85A and K792A at interaction interfaces (Fig. 6E). These results indicated that filament clustering may be essential for the anti-phage function of *Ef*Avs5.

**Fig 6.**
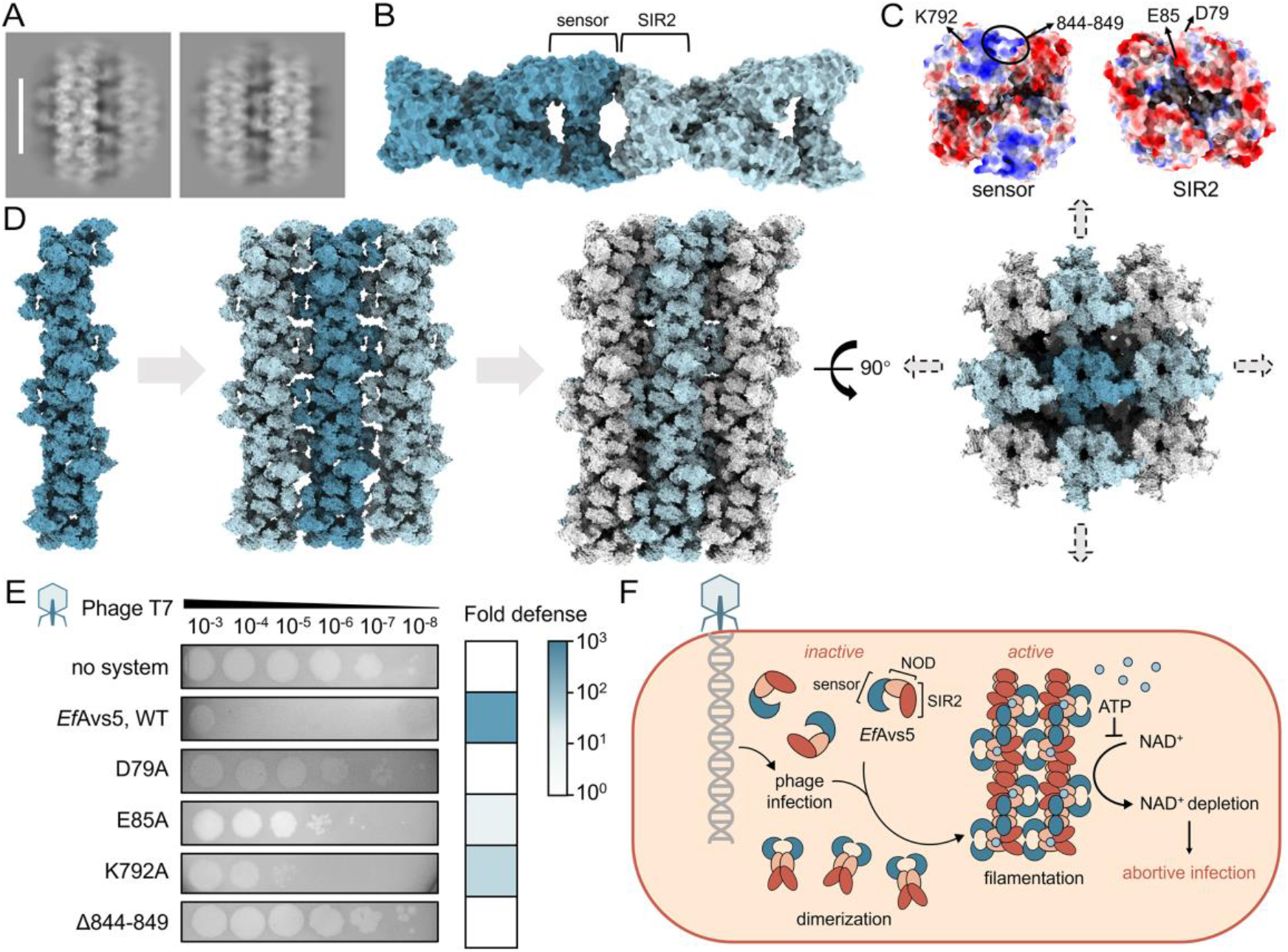
Filamentous cluster formation mediates *Ef*Avs5 anti-phage activity. (**A**) 2D class averages of the particle stacks, showing the bundled filament assembly. The scale bar represents 130 Å. (**B**) Interacting *Ef*Avs5 dimers in adjacent filaments. (**C**) The electrostatic potential interface of C-terminal sensor and N-terminal SIR2 domain. (**D**) A schematic diagram of the filament cluster assembly of *Ef*Avs5. (**E**) Phage plaque assay of *E. coli* expressing *Ef*Avs5 variants with mutations at the clustering interfaces. Data are representative images of n=3 biological replicates. (**F**) Proposed model of *Ef*Avs5 immune mechanism. Without phage infection, *Ef*Avs5 remains inactive as monomer. Phage infection activates *Ef*Avs5, forming filament bundles where SIR2 enable rapid NAD^+^ degradation and aborting phage infection.

## Discussion

In this study, we determined the cryo-EM structures of the *Ef*Avs5 and found that *Ef*Avs5 assembles into bundled filaments for NADase activity and anti-phage defense. We propose the following model for *Ef*Avs5 (Fig. 6F). Without phage infection, *Ef*Avs5 adopts an inactive, monomeric conformation in cells. Phage infection triggers *Ef*Avs5 to form filament bundles, in which SIR2 adopts a dimeric active state. This active state enables *Ef*Avs5 to rapidly degrade NAD^+^, leading to an abortive infection that prevents propagation of phage through a population of cells.

The *Ef*Avs5 structure displays a filament assembly distinct from the characterized SIR2-containing defense systems and STAND proteins, underlining the universal yet diverse nature of filamentation as an antiviral mechanism across prokaryotic and eukaryotic immune systems. The *Ef*Avs5 filament is formed by *Ef*Avs5 dimers with a C2 symmetry arranged into a high-order complex, involving comprehensive SIR2-SIR2 and NBD-WHD interactions. In SIR2-APAZ/Ago, DSR2, SIR2-HerA and Thoeris system, SIR2 domains function as monomers, tetramers (dimer of dimers), dodecamers (hexamer of dimers) and filament (spiral of tetramers), respectively^7-10^. Compared with the SIR2-based filamentation in ThsA, the NOD and SIR2 domain work together to facilitate filament formation in *Ef*Avs5, highlighting their independent yet complementary roles in the assembly process. Moreover, to the best of our knowledge, *Ef*Avs5 represents the first STAND family protein reported to assemble into functional filaments. The activation of STAND family proteins typically involves oligomerization, which includes tetramers (*Se*Avs3, *Ec*Avs4, and plant TIR-NLRs), pentamers (plant ZAR1), heptamers (animal apoptosome Apaf-1) or octamers (Drosophila dark apoptosome)^4,21,23-25^. Notably, *Solanum lycopersicum* NRC2 (*Sl*NRC2) was recently found to form filaments at elevated concentrations, but adopt an inactive conformation to avoid self-activation. Distinct from *Ef*Avs5, the *Sl*NRC2 filament consists of three identical protofilaments, which contain copies of tetramers^26^.

The activation observed in this study appears to be correlated with the accumulation of *Ef*Avs5 proteins, mirroring previous findings in plants where concentration-dependent hypersensitive response cell death was mediated by the plant NLR protein^27,28^. Considering its genomic localization on a PICI, intracellular expression of *Ef*Avs5 may be tightly regulated to avoid unnecessary activity. The phage signals activating *Ef*Avs5 remain unclear. We speculated that the activation of *Ef*Avs5 upon phage infection may involve binding of an as-yet-unidentified protein or compound; alternatively, it may be a response to changes in substrate concentration. Firstly, *Ef*Avs5 may interact with either phage-derived proteins, akin to the DSR2 system^9^, or host-derived proteins, such as that involved in the bNACHT25 system^29^, leading to the activation of filament formation and enzymatic activity. However, our attempt to identify potential activators through screening of escape phages was unsuccessful. Secondly, as suggested by the finding that the binding of the TIR substrates NAD^+^ and ATP induces phase separation and activation of TIR *in vitro*^*30*^, one of the potential activation mechanisms of *Ef*Avs5 may be the substrate-based response.

Above all, our structural and biochemical findings offer insights into the molecular mechanism of Avs5 bacterial immune system, highlighting the conservation of filamentation as a key regulatory mechanism governing antiviral defense across prokaryotes and metazoans.

### Limitations of the study

Our findings elucidate the fundamental mechanisms underlying *Ef*Avs5 function, yet the activation signals associated with phage infection remain elusive and warrant further investigation. Apart from biochemical assays on mutants, our results lack direct evidence of *Ef*Avs5 forming fibrillar clusters *in vivo*. Additionally, we propose that ATP intricately modulates *Ef*Avs5 activity, but the underlying molecular mechanisms require detailed clarification.

## Acknowledgements

This work was supported by National Natural Science Foundation of China (32370071, 32070040), National Key Research and Development Program of China (2020YFA0907900, 2019YFA0905400). We thank the staff members of the Electron Microscopy System at the National Facility for Protein Science in Shanghai (NFPS), Shanghai Advanced Research Institute, Chinese Academy of Sciences, China for providing technical support and assistance in data collection. We thank Jing Liu, Xinqiu Guo and Mengyu Yan at the Instrument Analysis Center (IAC) of Shanghai Jiao Tong University for providing assistance in data collection.

## Author contributions

J.Z. and Y.W. designed the experiments and analyzed data.

Y.W. and Y.T. performed biochemical assays.

Y.W. and X.Y. conducted cryo-EM data collection.

Y.W. conducted image processing, atomic model, building and refinement and structural analyses.

Y.W. and Y.F performed bioinformatics analysis.

Y.W. and J.Z. wrote the manuscript.

## Data availability

The atomic coordinates have been deposited in the Protein Data Bank under accession codes 9JAP. The corresponding maps have been deposited in the Electron Microscopy Data Bank under the accession number EMD-61299. Original data for biochemical assays and uncropped gels were deposited to Figshare.

## Supplemental information

Document S1. Figures S1–S10 and Table S1-S2

## Methods

### Plasmid construction

DNA fragments encoding *Ef*Avs5 from *Escherichia fergusonii* PICI EfCIRHB19-C05 (QML19490.1) were synthesized and inserted into the expressing plasmids pET28a with N-terminal His_6_-tag by Universe Gene Technology (Tianjin). The DNA fragments encoding *Ef*Avs5 was subcloned into the pBAD33 vector for phage infection assays. Plasmids carrying *Ef*Avs5 mutants were constructed using site-directed mutagenesis. Primers used are listed in Table S2.

### Protein expression and purification

The vector pET28a-*Ef*Avs5 was transformed into *E. coli* BL21(DE3). Induction of expression was achieved by adding 0.3 mM isopropyl-β-D-thiogalactopyranoside (IPTG) and incubated for 16 h at 16°C and 220 rpm. The cells were then harvested and resuspended in buffer A (50 mM Tris, pH 8.0, 500 mM NaCl, 5 mM MgCl_2_, 10% glycerol, 1 mM β-mercaptoethanol and 5 mM imidazole). After purification by nickel-NTA, the eluate was further loaded onto a size exclusion chromatography column (SEC) (Superose 6, Cytiva) equilibrated with buffer B (20 mM Tris, pH 8.0, 150 mM NaCl, 5 mM MgCl_2_, 5% glycerol) and the purified proteins were stored at −80 °C.

### Plaque assays

*E. coli* MG1655 possessing pBAD33-*Ef*Avs5, its mutants or an empty vector(pBAD33) were grown at 37°C in LB medium. When the OD_600_ reached 0.4, cells were induced with 0.5% L-arabinose for 1 hour. Then, 150 μL of the culture was mixed with 3 mL LB containing 0.8% low-melting agarose (with 0.5% L-arabinose added), and the mixture was poured into LB agar base layer in the 9 cm petri dish. Ten-fold dilutions of high-titers(>10^8^pfu/mL) of phage T7 were spotted onto the agar and incubated overnight at 37 °C. The next day, plates were photographed with blue-white light reflection in the dark box.

### Phage-infection in liquid medium

*E. coli* MG1655 possessing pBAD33-*Ef*Avs5, its mutant(N141A) or an empty vector(pBAD33) were grown until the OD_600_ reached 0.4. These cultures were then induced with 0.5% L-arabinose for 1 hour, and diluted in LB containing 0.5% L-arabinose to an OD of ∼0.1. Phage T7 was added to the culture at final MOI of 0.02, 0.2, 2, and 10. Subsequently, 200 μL of the diluted culture was placed into the wells of a 96-well plate. During shaking, the OD_600_ was measured every 5 minutes at 37°C for 6 hours.

### Cell survival assays

Overnight cultures of *E. coli* MG1655, either containing the plasmid pBAD33-*Ef*Avs5 or an empty vector, were diluted 1:100 into fresh LB medium. These cultures were grown at 37°C with the addition of 0.5% L-arabinose until they reached an OD_600_ of 0.4. Subsequently, the cells were harvested by centrifugation, washed with LB, and resuspended to an OD_600_ of 0.2. To each sample, phage T7 was added at an MOI of 5, while control samples were left without phage addition. Following a 20-minute adsorption period, serial 10-fold dilutions of each sample were plated onto LB agar, and the plates were incubated overnight at 37°C. The cell survival rate was determined by comparing the CFU obtained from the samples with phage T7 addition to the CFU obtained from the control samples without phage, expressed as a percentage.

### Phage burst size measurements

Overnight cultures of *E. coli* MG1655 harboring either the pBAD33-*Ef*Avs5 plasmid or an empty vector were diluted 1:100 in LB medium supplemented with 0.5% L-arabinose. These diluted cultures were then grown at 37°C until they reached an OD_600_ of 0.4. Subsequently, T7 phages were introduced to the cultures at an MOI of 0.1, and the infection process was allowed to proceed at 37°C. To establish a baseline for the initial phage titer, an equal volume of phage was added to LB media and used as the reference for time 0 of infection. After infection for 20, 40, and 60 minutes at 37°C, which represent approximately one, two, and three cycles of T7 phage replication, respectively, 0.5 mL samples of the culture were withdrawn. These samples were centrifuged at 5000 rpm for 7 minutes and the supernatants were filtered through a 0.22 μm filter. The titer of the T7 phages present in the filtered supernatants was determined by a plaque assay using *E. coli* MG1655 as the host.

### NAD(H) degradation measurements

NAD(H) concentrations were measured using the Innochem Coenzyme I NAD(H) Content Assay Kit (WST colorimetry), following the manufacturer’s instructions. Briefly, overnight cultures were diluted 1:100 in 50 mL LB with 0.5% L-arabinose, grown to OD_600_ of 0.4, and infected with T7 phage at an MOI of 2. At indicated times, 1 mL cultures were centrifuged, resuspended in acidic extract, sonicated, boiled, rapid cooled, and centrifuged. Supernatant was neutralized with alkaline extract. 50 μL of the neutralized supernatant was combined sequentially with 250 μL of Reagent 1, 75 μL of Reagent 2, 150 μL of Reagent 3, and 35 μL of Reagent 4 in the determination tube. After a 1-hour dark reaction, Reagent 5 was added to the mixture. In the control tube, Reagent 5 was added first, followed by the supernatant, maintaining identical steps thereafter. Absorbance was measured at 450 nm, and NAD(H) levels were calculated using a standard curve and normalized to the amount of NAD(H) present in an equivalent volume of sample with an OD_600_ of 0.1 to allow for accurate comparisons between samples.

### Negative staining analysis

The *Ef*Avs5 sample was directly applied to a glow-discharged 400-mesh Cu grid (Beijing ZhongJingKeYi Technology) for 30 s. After side blotting, the grid was immediately stained with 2% uranyl formate and then blotted again from the side. Staining was repeated twice with a 30 s incubation with uranyl formate in the final staining step. EM images were collected on a Talos F200C G2 at a nominal magnification of 52,000× and at a defocus of about 5 µm.

### Cryo-EM grid preparation and data acquisition

3.5 μL of the *Ef*Avs5 samples were added to freshly glow-discharged Quantifoil R1.2/1.3 Cu 300 mesh grids. In a Vitrobot (FEI, Inc.), grids were plunge-frozen in liquid ethane after being blotted for 2 s at 16°C with 100% chamber humidity. The grids were imaged using EPU on Titan Krios 300 kV microscopes with a K3 detector. Totally 4318 movies were collected under the defocus ranged from −0.8 to −1.6 μm, and the magnification was 81k in super-resolution mode. 32 frames per movie were collected with a total dose of 40 e^−^/Å^2^.

### Cryo-EM data processing and model building

The cryo-EM data was processed using CryoSPARC suite v4.5.3^31^. After motion correction, patch CTF estimation and manual exposure curation, 1425 movies were selected. The first set of particles was picked using a filament tracer, and after two-dimensional (2D) classification, good templates were selected for template picking. A total of 1,392,237 particles were picked using a template picker and extracted (box size 360 pixels) from 1,425 accepted micrographs (Fig. S4). Two rounds of 2D classification were conducted and 364,551 particles were picked and used for ab-initio reconstruction (three classes). The largest 3D class (228,053 particles, 62.6%) was selected and refined by homogeneous refinement, non-uniform refinement and CTF refinement to give the final map (global resolution, 3.44 Å). Helical parameters (helical twist and helical rise) were determined by helical refinement job. The initial model of *Ef*Avs5 wes generated by AlphaFold^32^. The model of ATP was built in Coot^33^. The models of *Ef*Avs5 were fitted into the cryo-EM density maps using ChimeraX^34^. The model was refined in Coot and Phenix with secondary structure, rotamer and Ramachandran restraints^35^. The Map versus Model FSCs was generated by Phenix (Fig. S4). The statistics of cryo-EM data processing and refinement were listed in Table S1.

### *In vitro* NADase activity

The *Ef*Avs5 NADase activity was evaluated through an εNAD^+^-based fluorescence assay, wherein the enzymatic cleavage of the nicotinamide glycosidic bond in εNAD^+^ results in the formation of εADPR, which subsequently emits a fluorescent signal. In a 96-well black flat-bottom plate, 100 μL reaction mixtures were prepared, each containing 50 μM of εNAD^+^ (Sigma catalogue No. N2630) and 0.5 μM *Ef*Avs5 proteins in reaction buffer (20 mM Tris, pH 8.0, 150 mM NaCl, and 5% glycerol, with and without the addition of nucleotides). A mixture containing only the reaction buffer and εNAD^+^ was used as a control. The reactions were then loaded into a BioTek Synergy H1 microplate reader, and fluorescence intensities were recorded at 37°C every 20 seconds for 15 minutes, employing excitation and emission wavelengths of 310 nm and 410 nm, respectively. All experiments were conducted in triplicate to ensure reproducibility.

### ATPase activity

ATPase activities were evaluated by using Malachite Green Phosphate Detection Kit (Beyotime). Each 200 μL reaction mixture consisted of 5μM *Ef*Avs5 proteins in a reaction buffer (20 mM Tris, pH 8.0, 150 mM NaCl, 5 mM MgCl_2_ and 0.5 mM ATP). A mixture without the addition of *Ef*Avs5 proteins served as a control. Following a 30-minute incubation at 37°C, 70 μL of color reagent was added and the mixture was further incubated for another 30 minutes. Subsequently, the absorbance at 620 nm was measured using a multi-detection microplate reader (Tecan Spark). Phosphate levels were then calculated using a standard curve generated with phosphate standards. All experiments were conducted in triplicate to ensure reproducibility.

### Phylogenetic analysis of Avs5 systems

The coding sequence of Avs5 was acquired from two sources: firstly, by utilizing the defense finder tool to scan through all bacterial and archaeal genomes housed in the IMG (Integrated Microbial Genomes & Microbiomes) database from November 2023, aiming to identify proteins that encode the Avs5 system^36^; secondly, by leveraging the Foldseek search server to discover proteins (leveraging AlphaFold/UniProt50, ensuring a coverage exceeding 80%) that exhibited a high degree of structural similarity to Avs5^37^. To eliminate redundancy, MMseqs was employed with the specified parameters ‘–min-seq-id 0.90’ and ‘-c 0.8’, resulting in 263 filtered sequences^38^. These sequences were subsequently aligned using MAFFT, with the parameters set to ‘--maxiterate 1000 –globalpair’ for optimal alignment^39^. Following alignment, trimAl was applied to refine the alignment by trimming unnecessary regions^40^. The phylogenetic tree was then constructed using IQ-TREE, incorporating the parameters ‘-nstop 500 -bb 1000 -m LG+F+I+G4’ for enhanced accuracy and robustness^41^. Finally, iTOL was utilized for the visualization and annotation of the phylogenetic tree^42^.

