## Supplemental Data 1 for "Filamentation activates bacterial NLR-like antiviral protein"

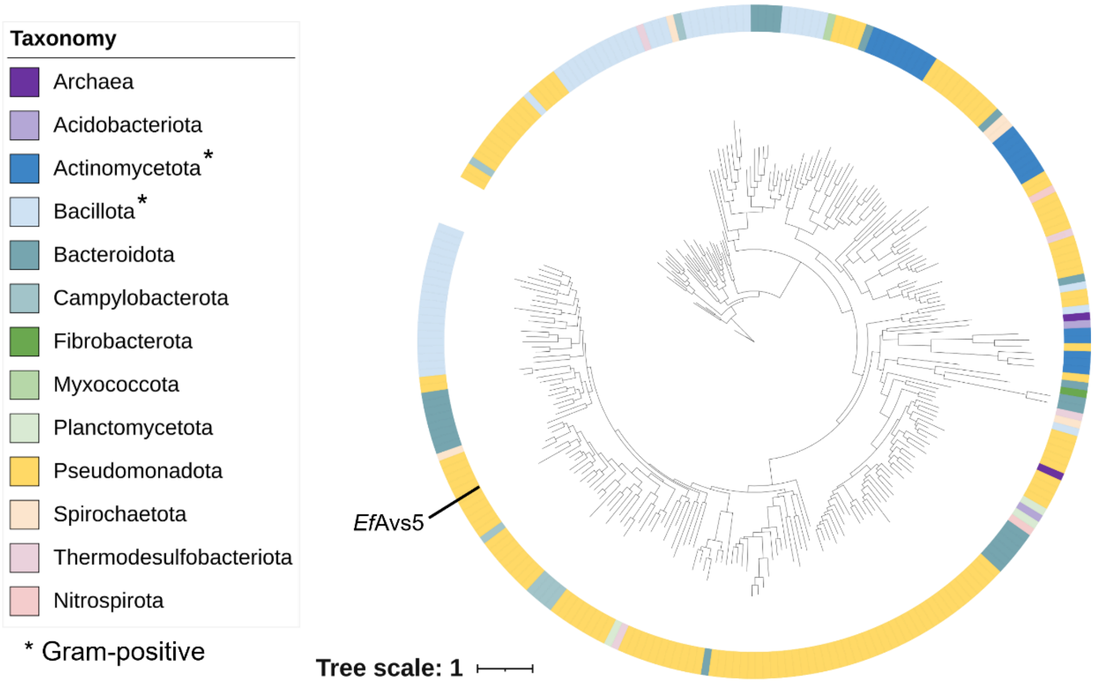


**Figure S1.** Avs5 phylogenetic tree (n=263).

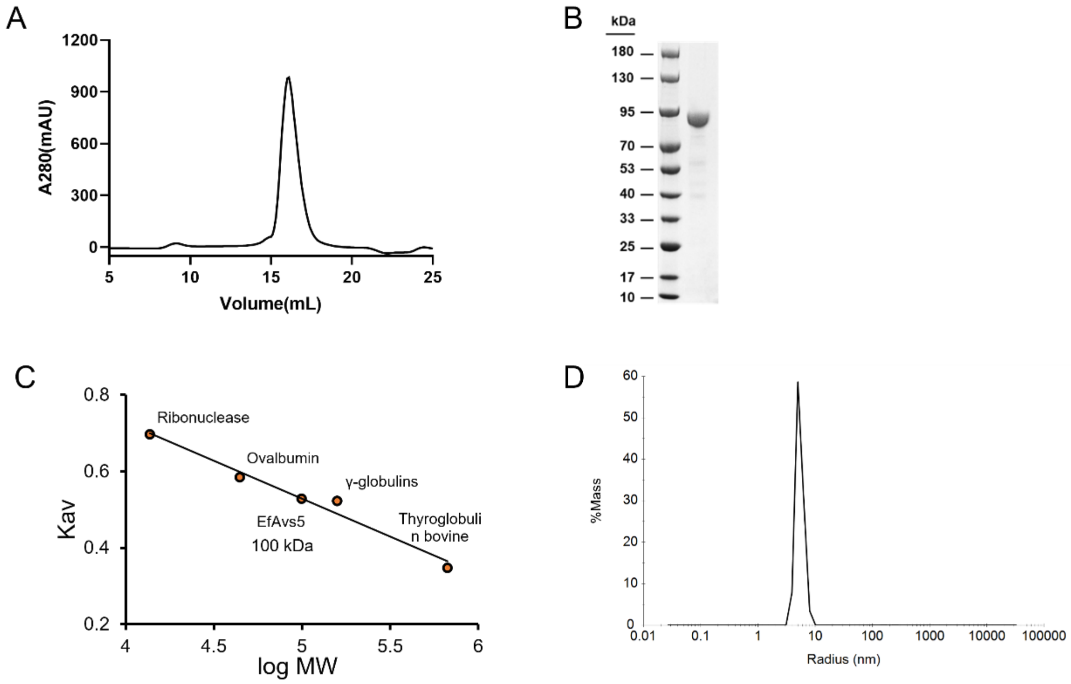


**Figure S2.** The freshly isolated *Ef*Avs5 exists as soluble monomer. (A) Size exclusion chromatography analysis of *Ef*Avs5 using Superose 6 Increase column. (B)SDS-PAGE of *Ef*Avs5. (C) Standard curve generated using Superose 6 Increase column to determine the molecular weight of *Ef*Avs5 in the solution. The molecular weight of *Ef*Avs5 was determined to be approximately 100 kDa. (D) Dynamic light scattering (DLS) analysis of *Ef*Avs5, showing a particle radius of approximately 5.6 nm. Data represents the calculated average from five separate measurements.

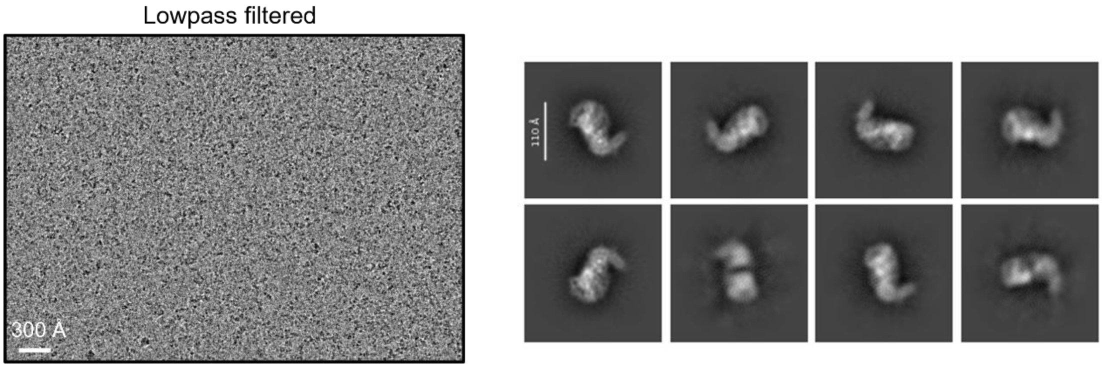


**Figure S3**. Representative cryo-EM micrograph (left) and 2D classes (right) of the monomer *Ef*Avs5. The scale bar represents 300 Å in the cryo-EM images and 110 Å in the 2D images.

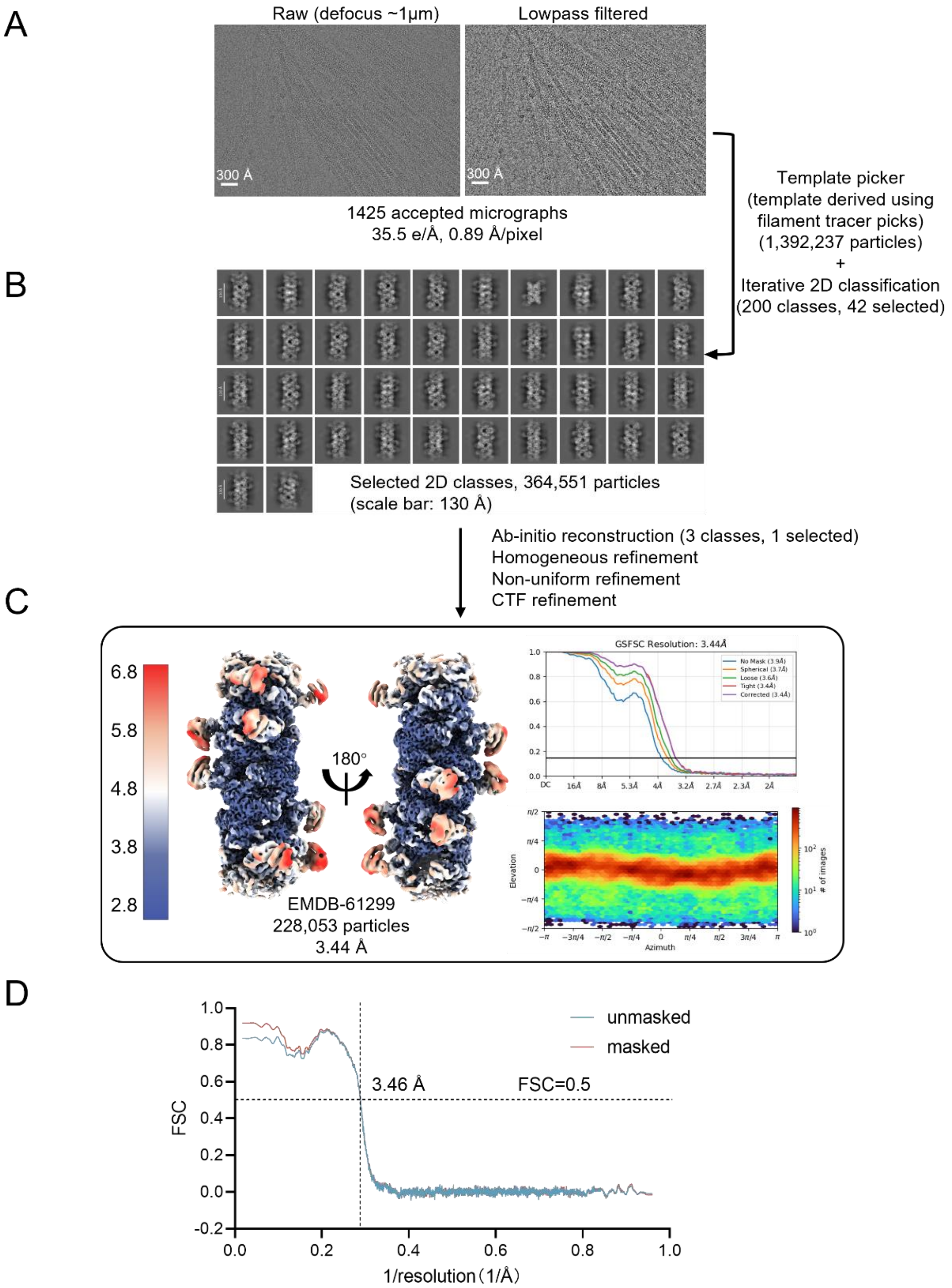


**Figure S4**. Cryo-EM data processing and validation for *Ef*Avs5. (A) Representative cryo-EM micrograph for the final reconstruction. The scale bar represents 300 Å. (B) 2D class averages of the particle stacks, showing the filament assembly. The scale bar represents 130 Å.(C) The final cryo-EM map of the *Ef*Avs5 was reconstructed using a total of 228,053 single particles and refined to a nominal resolution of 3.44 Å. (D) Validation of cryo-EM structural models. Map vs. model FSCs was generated by Phenix (Version 1.21.1). The model-map resolution for the atomic model and cryo-EM map at FSC = 0.5 cutoff was indicated in the figure and reported in Table S1.

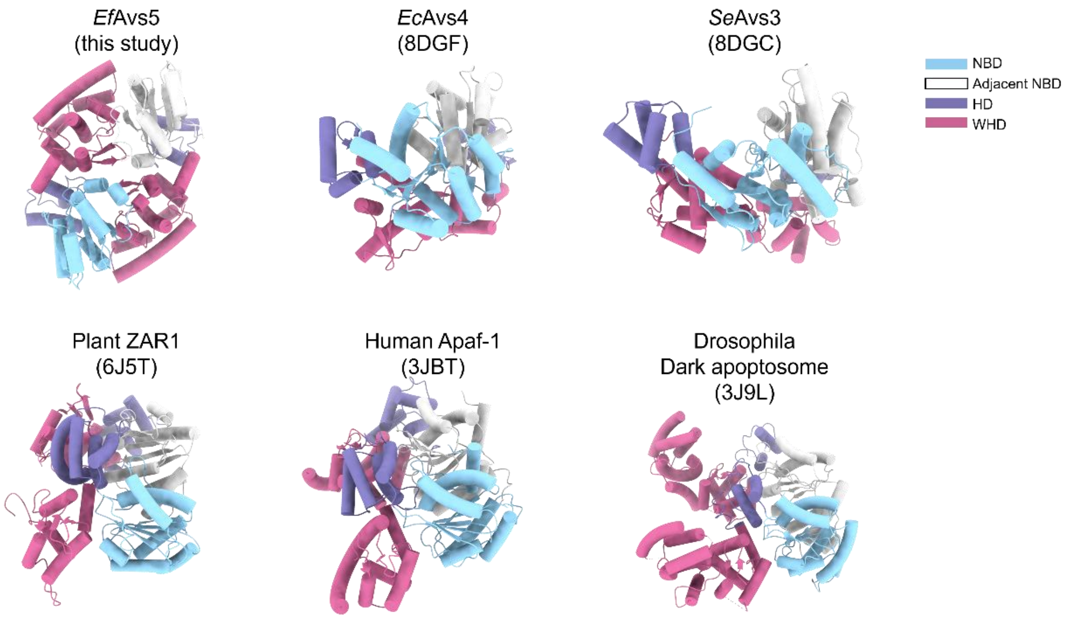


**Figure S5**. Comparison of the activated STAND structures, showing different relative positions of the adjacent NOD. NBD; nucleotide-binding domain. HD; helical domain. WHD; winged-helix domain.

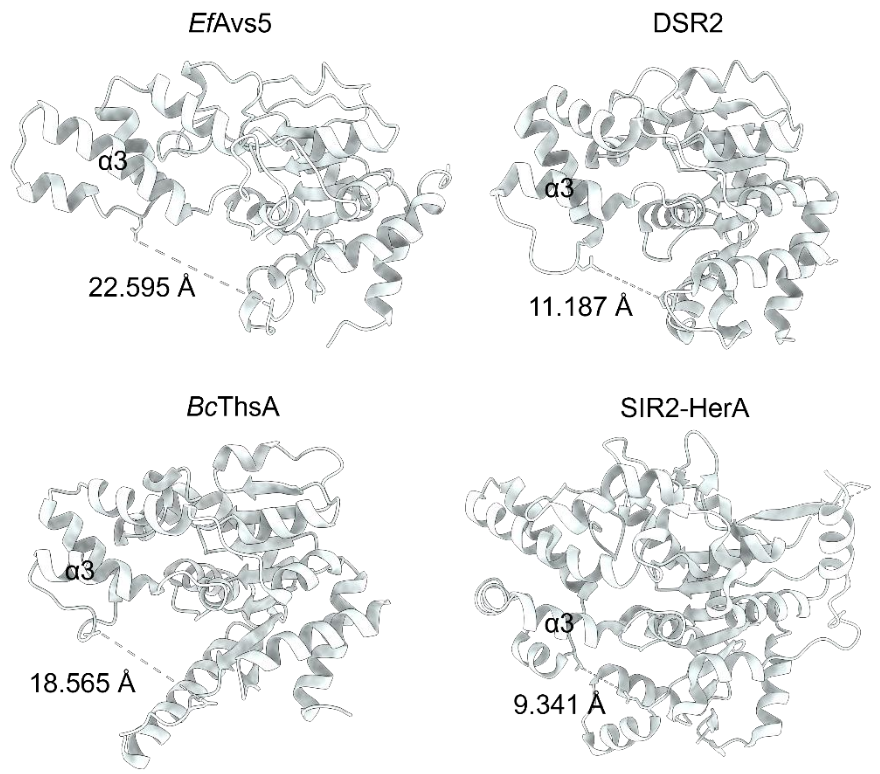


**Figure S6**. Structure comparison of SIR2 domain in Avs5, DSR2, Thoeris and the SIR2-HerA defense systems.

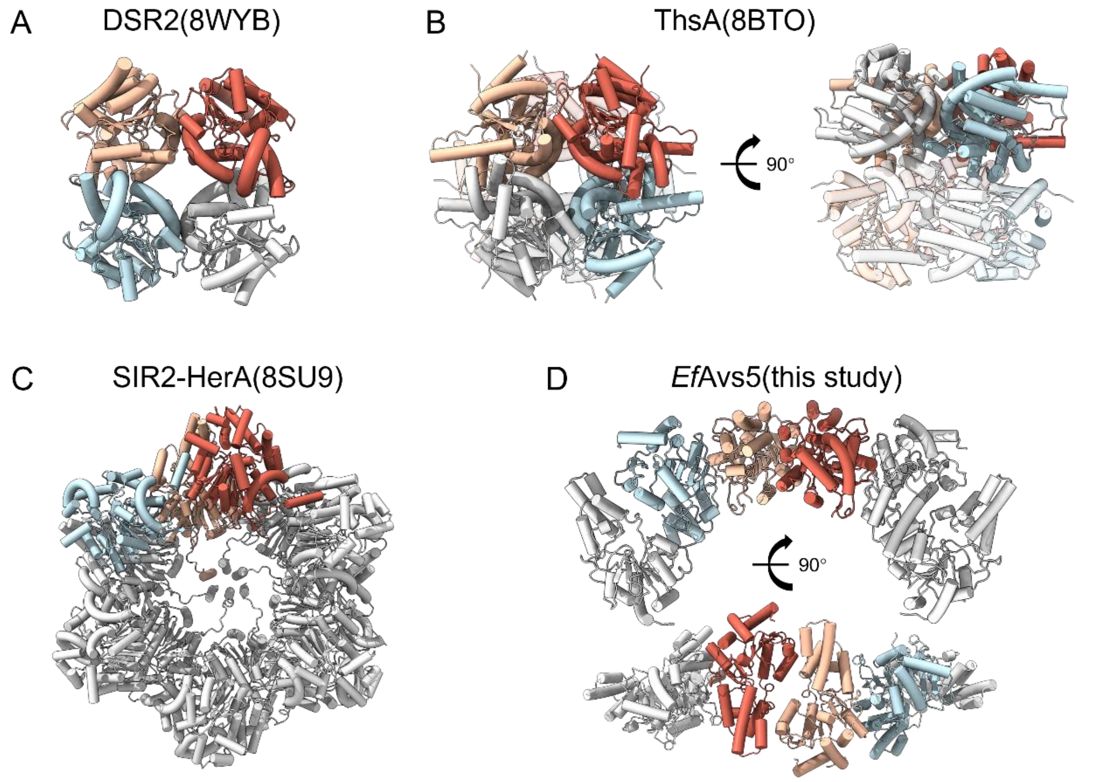


**Figure S7.** Comparison of the SIR2-assembly mechanisms in DSR2, Thoeris, SIR2-HerA and Avs5 defense systems.

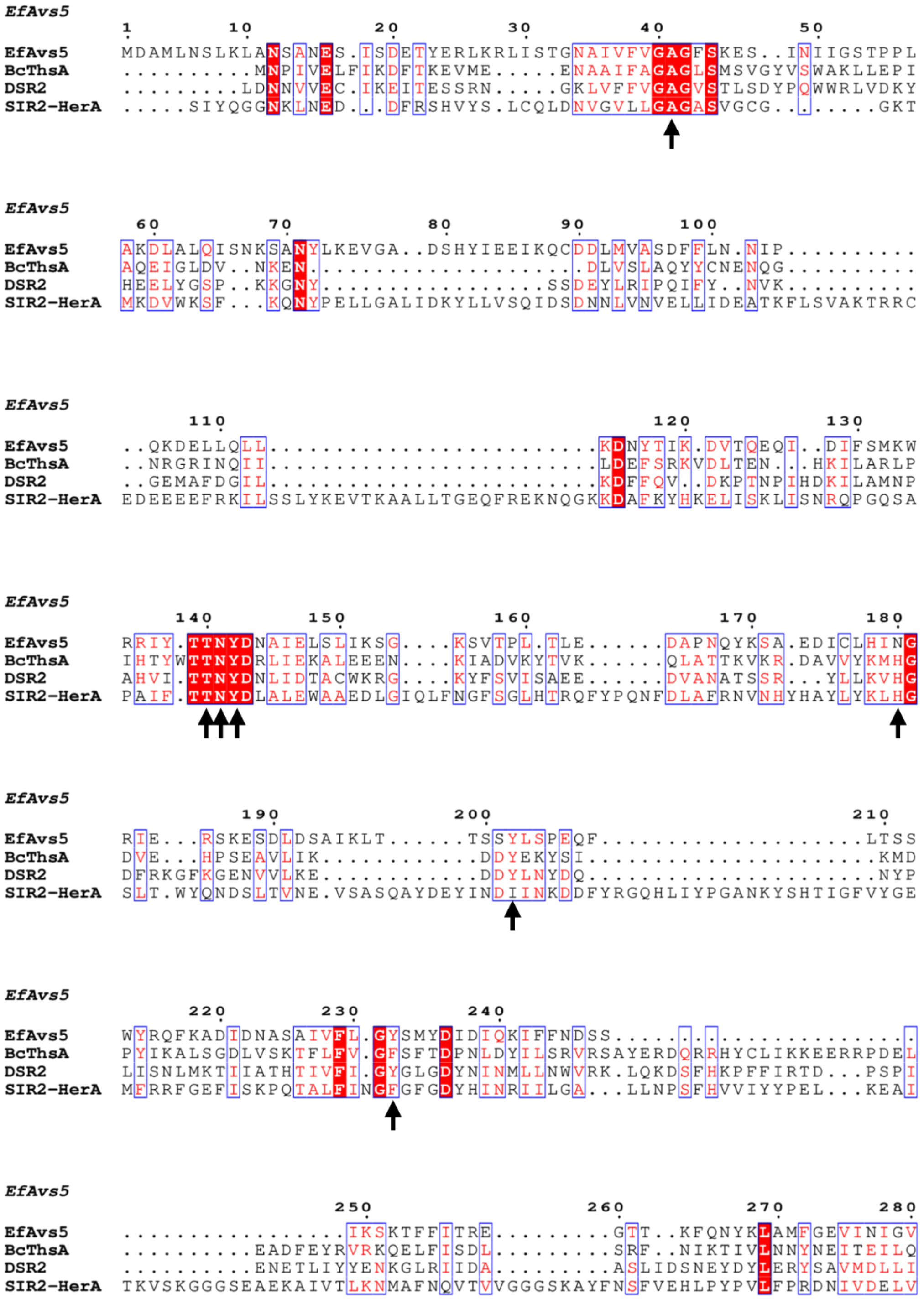


**Figure S8.** Sequence comparison of SIR2 domain in Avs5, DSR2, Thoeris and SIR2-HerA defense systems. The residues in the catalytic pocket make hydrogen bonds with $NAD^+$ are indicated by arrows.

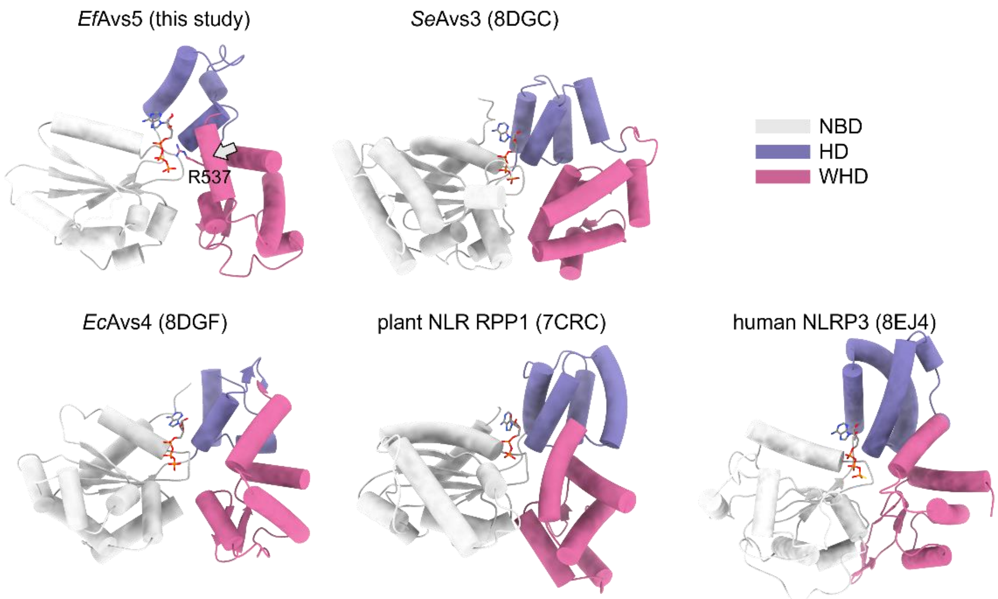


**Figure S9**. Comparison of the NOD structure of activated STAND proteins. The main difference between Avs5 and other STANDs is indicated by a grey arrow.

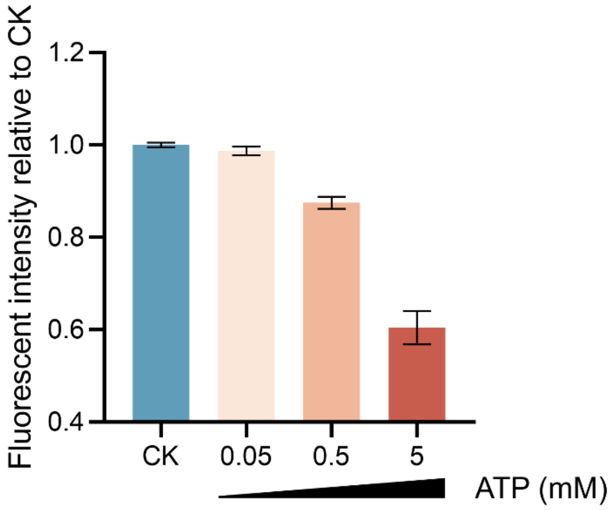


**Figure S10.** Effects of ATP on NADase activity of the *Ef*Avs5 complex. Data represent the mean of three replicates and error bars represent the SEM.

**Table S1**. Cryo-EM data collection, refinement, and validation statistics.

| | *Ef*Avs5 (EMD-61299) (PDB 9JAP) |
|---|---|
| **Data collection** | |
| Magnification | 81,000 |
| Voltage (kV) | 300 |
| Electron exposure($e^-$/Å$^2$) | 40 |
| Defocus Range (μm) | 0.8-1.6 |
| Pixel Size (Å) | 0.89 |
| Initial particle images (no.) | 1,392,237 |
| Final particle images (no.) | 228,053 |
| Map resolution (Å) | 3.44 |
| FSC threshold | 0.143 |
| Map resolution range (Å) | 2.8-6.8 |
| **Refinement** | |
| Model resolution (Å) | 3.46 |
| FSC threshold | 0.5 |
| CCMask | 0.84 |
| Model composition | |
| Non-hydrogen atoms | 109208 |
| Protein residues | 6664 |
| Nucleotides | 0 |
| Ligands | 16 |
| B-factors (Å$^2$) | |
| Protein | 82.03 |
| Nucleotides | --- |
| Ligands | 34.38 |
| RMS deviations | |
| Bond length (Å) | 0.005 |
| Bond angle (°) | 1.028 |
| **Validation** | |
| MolProbity score | 1.63 |
| Clash score | 5.69 |
| Rotamer outliers (%) | 1.51 |
| Ramachandran Outliers (%) | |
| Outliers | 0 |
| Allowed | 3.04 |
| Favored | 96.95 |

**Table S2**. List of primer sequences used in this study.

| Primer Name | Sequence (5' to 3') |
|---|---|
| AVS5 D79A F | GAAGGAAGTTGGTGCAGCAAGCCATTATATTGAA |
| AVS5 D79A R | TGCTGCACCAACTTCCTTCAGAT |
| AVS5 E85A F | TAGCCATTATATTGAAGCAATCAAGCAGTGCGAC |
| AVS5 E85A R | GCTTCAATATAATGGCTATCTGCAC |
| AVS5 N141A F | CCGTATCTATACCACCGCCTATGATAATGCCATC |
| AVS5 N141A R | GGCGGTGGTATAGATACGGCG |
| AVS5 D164A F | CCCGCTGACCCTGGAAGCCGCCCCTAATCAGTAT |
| AVS5 D164A R | GGCTTCCAGGGTCAGCGGGGTCACACT |
| AVS5 R185A F | TAATGGTCGCATTGAAGCCAGTAAAGAAAGTGAT |
| AVS5 R185A R | GCTTCAATGCGACCATTAATATGCAGG |
| AVS5 Y202A F | GCTGACCACCAGTAGTGCACTGAGTCCGGAACAG |
| AVS5 Y202A R | TGCACTACTGGTGGTCAGCTTAATTG |
| AVS5 K241A F | TGACATTGACATTCAGGCAATCTTCTTCAACGA |
| AVS5 K241A R | CTGAATGTCAATGTCATACATAC |
| AVS5 T262A F | ACCCGCGAAGGTACCGCAAAATTTCAGAATTAT |
| AVS5 T262A R | GGTACCTTCGCGGGTGATGAAA |
| AVS5 I303AE307A F | TAAGGAAGTGGGCCTGGCAAATAGTCTGGCACTGTATAC |
| AVS5 I303AE307A R | TGCCAGGCCCACTTCCTTATCCTGATGA |
| AVS5 H362A F | GAGCAAAATTATTGAAGCCATCGAGACCGATAAC |
| AVS5 H362A R | GCTTCAATAATTTTGCTCACTTCTTCA |
| AVS5 K379A F | AGCGATCTGGGCAATGGTGCCAGTATTATGACCCGT |
| AVS5 K379A R | GGCACCATTGCCCAGATCGCTGGCAATC |
| AVS5 E404A F | TTATTACCTGTACAACGCCTTCAGCTTCAGCAAA |
| AVS5 E404A R | GGCGTTGTACAGGTAATAAAAACACAG |
| AVS5 D426A F | AATTGTTATTTTCATCGCCGATTACAGCAATTGC |
| AVS5 D426A R | GCGATGAAAATAACAATTTTCTGGCC |
| AVS5 K461A F | TGGCTATGAAAATACCGCACAGCACCTGCTGACC |
| AVS5 K461A R | GCGGTATTTTCATAGCCAAAATGACGG |
| AVS5 K552A F | GAATGATAAAGAATATGCCGAAACCGTGTTCGCC |
| AVS5 K552A R | GGCATATTCTTTATCATTCAGGGCCAGG |
| AVS5 H622AK623A F | TGCCAATATTTTTGAGGCAGCATACGTGGTTAACCAG |
| AVS5 H622AK623A R | TGCCTCAAAAATATTGGCAATCAGAAAA |
| AVS5 Δsensor-F | TGAAAAGCTGCTGCCGTAACAGCGTCGTGTTGAA |

| | |
|---|---|
| AVS5 Δsensor-R | TTACGGCAGCAGCTTTTCAATAATACT |
| AVS5 K792A F | CCCGACCTTTAACGCCGCACAGAAAGTGTTTTA |
| AVS5 K792A F | GCGGCGTTAAAGGTCGGGTACACTTTT |
| AVS5 Δ844-849 F | AAAATATCCAGGAAAACTAACGCCCGAAAGGTAAA |
| AVS5 Δ844-849 R | TTAGTTTTCCTGGATATTTTCCACAATC |
